# Structural characterization of stochastic detailed balance in chemical reaction networks

**DOI:** 10.64898/2026.07.24.740452

**Authors:** Shangbin Ma, Youming Li

## Abstract

The global potential of a chemical reaction network has many applications and is closely related to the stochastic detailed balance. However, many fundamental questions concerning stochastic detailed balance remain unresolved, such as whether it depends on the system volume and how to construct new systems that satisfy it. In this paper, we show that stochastic detailed balance may depend on the system volume. We therefore introduce four types of stochastic detailed balance according to their dependence on volume and rate constants, and systematically investigate the relationships among them. Our results distinguish detailed balance arising from particular choices of volume and parameters from that enforced by network structure, and identify conditions under which detailed balance at one volume extends to all volumes. We further obtain a class of networks satisfying stochastic detailed balance for every volume and every positive choice of rate constants, and construct new systems whose global potentials exhibit double-well structures.

## 1 Introduction

Chemical reaction networks provide a unified framework for modeling interacting systems at both deterministic and stochastic levels, with applications including cell biology, epidemiology, ecology, sociology, and game theory [1, 2]. For a well-mixed system with a large number of molecules, fluctuations can be neglected and molecular concentrations are typically described by ordinary differential equations (ODEs). For systems having low molecular numbers, such as gene expression systems in which DNA and mRNA molecules are present in very low numbers [3, 4], the fluctuations are no longer negligible, and a suitable model is continuous-time Markov chain defined on nonnegative integer lattice. The continuous deterministic and discrete stochastic descriptions are linked by Kurtz’s law of large numbers: after concentration scaling, the stochastic trajectories converge to their deterministic counterparts on compact time intervals as the system size (or volume) tends to infinity [5–7]. Since the continuous deterministic model can be recovered as the large-system limit of the discrete stochastic model, the latter “is not an alternative to the deterministic kinetics, but a more complete kinetic description that is capable of modelling reactions with and without fluctuation” [8].

Analyzing continuous deterministic ODE models is generally simpler than solving the discrete stochastic counterparts, primarily due to the computational intractability arising from state-space explosion for the latter [9]. However, at small volumes, intrinsic noise becomes important, and the ODE predictions can differ substantially from those of the discrete stochastic model. This discrepancy cannot generally be captured even by standard Fokker-Planck or linear-noise approximations [10, 11]. These limitations motivate direct analysis or simulation of the discrete model, without relying on approximations.

Detailed balance provides an important structure for analyzing discrete stochastic models. The associated global potential, which plays a central role in statistical physics, further connects the discrete stochastic and continuous deterministic descriptions. In particular, when the system size is large, it can be used to characterize rare events in the stochastic model. However, the detailed balance in the deterministic and stochastic descriptions of chemical reaction networks differs in a fundamental way. In the deterministic framework, detailed balance requires the forward and backward rates of every reversible reaction pair to be equal at a positive steady state [12, 13]; for mass-action chemical systems, this is precisely the condition of chemical equilibrium [14]. In the stochastic Markov chain framework, detailed balance requires every transition between two states to be exactly balanced by the reverse transition under a stationary measure. It has been shown that deterministic detailed balance always implies stochastic detailed balance [15]. When multiple reactions share the same reaction vector, however, the converse may fail. In particular, an orthogonality condition has been proposed to produce systems satisfying stochastic detailed balance without satisfying deterministic detailed balance [16]. More importantly, such systems can exhibit double-well global potentials, in contrast to the single-well structure associated with deterministic detailed balance. This raises a natural question: apart from deterministic detailed balance and orthogonality condition, are there other mechanisms that lead to stochastic detailed balance?

Constructing a global potential requires stochastic detailed balance to hold for all system volumes [16]. This leads to the following question: does stochastic detailed balance in chemical reaction networks depend on the volume? We use counterexamples to show that it indeed may depend on volume. These counterexamples motivate us to define four types of stochastic detailed balance based on whether they are independent of volume and whether they are independent of parameters, and we analyze the relationships among these four types. Based on our discussion, we provide a class of systems satisfying universal stochastic detailed balance that is independent of both volume and parameters. We also provide more systems that satisfy stochastic detailed balance and have global potentials with double-well structure. This paper is organized as follows. In Section 2, we introduce models and preliminary results. In Section 3, we define the four types of stochastic detailed balance. In Section 4, we discuss the relationship between these four types and show how to construct new systems that satisfy stochastic detailed balance without deterministic detailed balance. We conclude in Section 5.

## 2 Model

### 2.1 Chemical reaction network

We consider a well-mixed chemical reaction system composed of a collection of chemical species *S* = *{S*_1_, …, *S*_*d*_*}* and a family of reactions

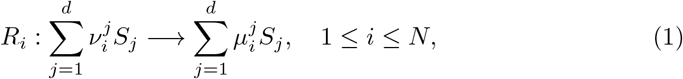

where 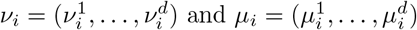 are the reactant and product complexes, respectively. The vector difference *µ*_*i*_ − *v*_*i*_ indicates the species change after the occurrence of the reaction and we call it the *reaction vector* of *R*_*i*_.

Let

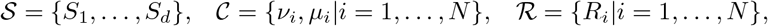

denote the collections of chemical species, complexes, and reactions, respectively. Then the ordered triple *{S, C, R}* is called a chemical reaction network [17].

According to standard conventions, a chemical reaction network is called *reversible* if for any reaction *R*_*i*_ : *v*_*i*_ → *µ*_*i*_ ∈ *R*, there exists a reverse reaction 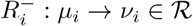. For any reversible pair, we uniquely classify *R*_*i*_ as a forward reaction if *v*_*i*_ *< µ*_*i*_ in the lexicographic order; otherwise, it is a backward reaction.

In complex chemical networks, such as those featuring parallel enzymatic and non-enzymatic pathways, different reactions may yield the same net change in species counts. This necessitates the following critical definition.

#### Definition 1

Two reactions are called **equivalent** if they have the same reaction vector.

Let *V* (*R*) = *{µ*_*i*_ − *v*_*i*_ | *R*_*i*_ is a forward reaction*}* denote the collection of distinct reaction vectors for forward reactions. The elements in *V* (*R*) are listed as *ω*_1_, …, *ω*_*r*_. For any *ω*_*p*_ ∈ *V* (*R*), we set

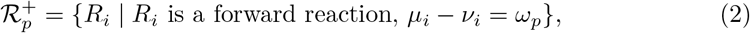

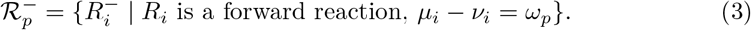

All reactions in 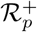 (together with their reverse reactions in 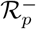) share the same reaction vector *ω*_*p*_ and are said to belong to the same *reaction family*. We relabel the elements in 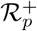 and 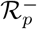 as

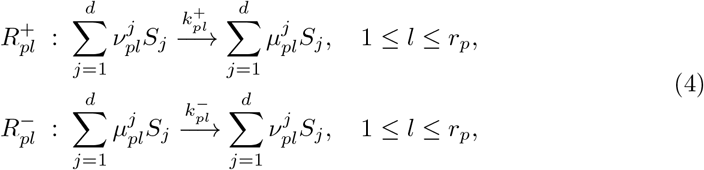

where *r*_*p*_ represents the number of forward reactions with the same reaction vector *ω*_*p*_.

### 2.2 Stochastic and deterministic models

For each 1 ≤ *j* ≤ *d*, let 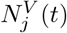 denote the number of molecules of chemical species *S*_*j*_ at time *t* with system size being *V*. Let 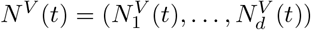, then the process {*N*^*V*^ (*t*) |*t* ≥0} can be modelled by a continuous-time Markov chain on *d*-dimensional nonnegative integer lattice with transition rates being [6]

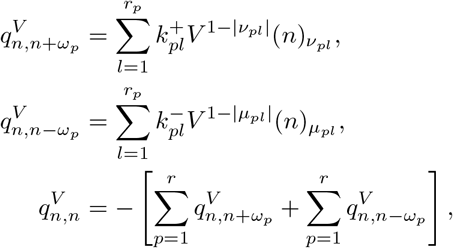

where, for *n* = (*n*_1_, …, *n*_*d*_) and *v* = (*v*^1^, …, *v*^*d*^),

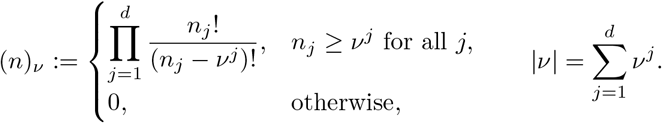

If *v* is a source complex of a reaction, then the number |*v* |is the *reaction order* of the reaction.

Clearly, the stochastic model evolves by jumps on the lattice 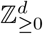. In general, not every state in this lattice is accessible from every other state. Since the reaction network is reversible and all rate constants are strictly positive, every possible jump can also be reversed. Consequently,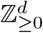 decomposes into disjoint closed irreducible communicating classes. The collection of all such classes is denoted by *C*. The class containing the initial state is the state space actually visited by the process.

We next introduce the continuous model, which is appropriate when molecular numbers are large. Let *x*_*j*_(*t*) be the macroscopic concentration of *S*_*j*_ at time *t*, then the evolution of *x*(*t*) = (*x*_1_(*t*), …, *x*_*d*_(*t*)) can be modelled by the following ordinary differential equations (ODEs):

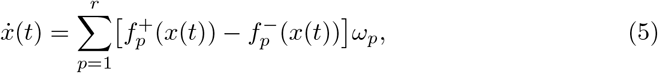

Where

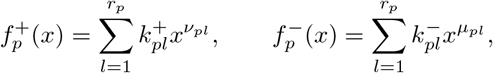

With 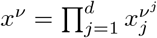. Kurtz’s law of large numbers states, under the usual hypotheses, that *N*^*V*^ (*t*)*/V* converges on compact time intervals to *x*(*t*) as *V*→ ∞ [5–7]. Beyond the law-of-large-numbers limit, the concentration process also satisfies a sample-path large-deviation principle, and Freidlin-Wentzell theory shows that the exponential cost of transitions between metastable states is governed by a quantity called quasi-potential. We refer interested readers to [16, 18, 19] for further details.

### 2.3 Detailed balance and global potential

For different models, detailed balance has different meanings: in the deterministic model, detailed balance means that at equilibrium each forward reaction flux is exactly balanced by its corresponding reverse reaction flux, so there is no net reaction flux along any reversible pair, a condition also called chemical equilibrium in chemistry. Mathematically, deterministic detailed balanced is equivalent to the existence of an equilibrium *c* satisfying

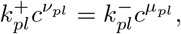

for every reversible reaction pair. In the stochastic model, detailed balance means that the probability flux from any state to any other state is exactly balanced by the reverse probability flux. The formal definition of stochastic detailed balance is as follows.

#### Definition 2

For fixed *V* > 0 and fixed positive rate constants, we say that the stochastic model satisfies **stochastic detailed balance** if, for every communicating class Γ ∈ *C*, there exists a positive measure

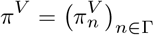

such that

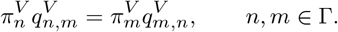

Here and below, only transitions between states in the same communicating class are considered. It has been proved that deterministic detailed balance implies stochastic detailed balance [15] at any *V*, which also implies local detailed balance [16] that guarantees the existence of global potential of chemical reaction systems. The following theorem presents the definition and some properties of the global potential.

#### Theorem 1 [16]

*Suppose that a reaction network satisfies stochastic detailed balance for any V with given parameters. Let* 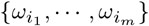 *be an arbitrary basis of* span(*V* (*R*)) *and let* 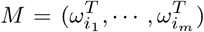 *be a d × m matrix, where m is the dimension of* span(*V* (*R*)). *Let* 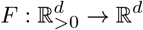 *be a vector field defined as*

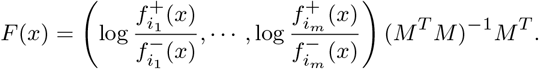

*Then the following statements hold:*

1. *The definition of the vector field F is independent of the choice of the basis of* span(*V* (*R*)). *In addition, for any* 1 ≤ *p* ≤ *r, we have*

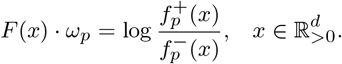
2. *The vector field F has a potential function* 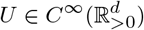, *namely*

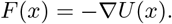
3. *The potential function U satisfies*

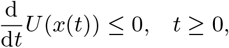

*where x* = *x*(*t*) *is the solution of the deterministic model* (5), *and the equality holds if and only if the deterministic model starts from any one of its equilibrium points*.

The global potential also has many properties in statistical physics. It solves the stationary Hamilton-Jacobi equation for the chemical large-deviation Hamiltonian and, inside a basin of attraction, agrees with the Freidlin–Wentzell quasi-potential [18] up to a constant. If the network satisfies deterministic detailed balance, *U* reduces to the classical free-energy function and is convex on the positive stoichiometric compatibility class [12]. The interesting case for us is therefore the systems that satisfy stochastic detailed balance but violate deterministic detailed balance. Such systems indeed exist, and can produce a double-well global potential. In this paper we will search for more such systems.

We now systematically analyze stochastic detailed balance. A simple method of verifying stochastic detailed balance is to use the Kolmogorov criterion [20], which states that a reaction network satisfies stochastic detailed balance at *V* if and only if the transition rates satisfy the Kolmogorov cycle condition. This means that every feasible cycle

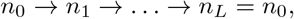

must satisfy

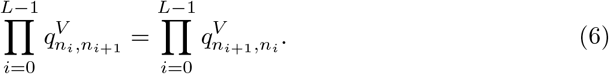

We now rewrite this condition explicitly. For each family *p*, define

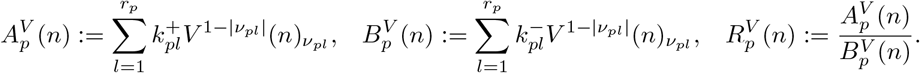

Since *µ*_*pl*_ = *v*_*pl*_ + *ω*_*p*_, we have

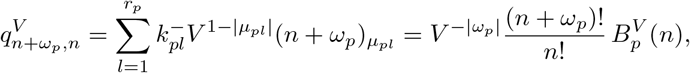

where 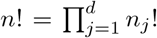 for *n* = (*n*_1_, …, *n*_*d*_). Therefore, whenever the transition *n* → *n* + *ω*_*p*_ is feasible, we have

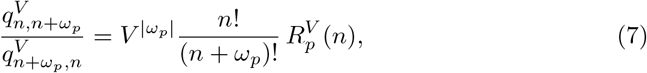

Where 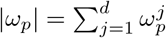. For a general cycle, write each transition as

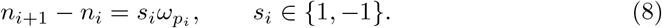

If the *i*-th step is a forward reaction, then *s*_*i*_ = 1 and its lower endpoint is the initial state *n*_*i*_. If it is a backward reaction, then *s*_*i*_ = −1 and its lower endpoint is the terminal state *n*_*i*+1_. We denote this lower endpoint by

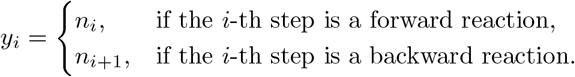

Substituting Eq. (7) into Eq. (6) gives the following equality that is equivalent to Kolmogorov’s cycle condition:

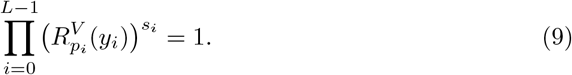

The above expression in terms of 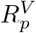 applies to all cycles. A standard cycle.decomposition result for lattice Markov chains states that every cycle can be decomposed into elementary cycles [21]. Consequently, if every elementary cycle satisfies the Kolmogorov cycle condition, then so does every cycle. In the stochastic chemical reaction networks considered here, these elementary cycles are of two types. The first is a parallelogram cycle generated by two linearly independent reaction vectors:

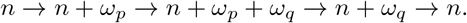

For this cycle, Eq. (9) gives

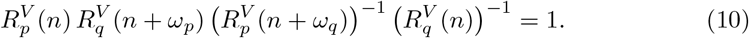

Equivalently,

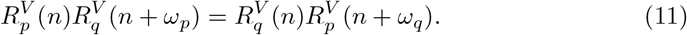

The second type of elementary cycle appears only when the reaction vectors are linearly dependent. Let *ξ* = (*ξ*_1_, …, *ξ*_*r*_) ∈ ℤ ^*r*^ *\ {*0*}* satisfy

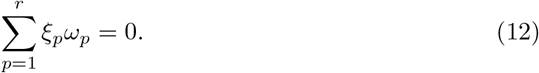

Choose an order of moves that uses *ξ*_*p*_ steps in the direction +*ω*_*p*_ if *ξ*_*p*_ > 0, and −*ξ*_*p*_ steps in the direction − *ω*_*p*_ if *ξ*_*p*_ *<* 0. Starting from a state sufficiently far from the boundary, these moves form a feasible linear-relation cycle.

#### Remark 1

We stress here that away from the boundary of the state space, parallelogram cycles and linear-relation cycles indeed constitute all elementary cycles. Near the boundary, however, some intermediate cycles in an algebraic decomposition may not be feasible, so a feasible cycle may not be decomposable into feasible elementary cycles of these two types. But this boundary issue does not affect the results. For example, the proof of Theorem 3 in [16] works directly with an arbitrary feasible cycle and establishes detailed balance along the entire cycle. For this reason, in this paper we only consider these two types of cycles for simplicity.

We now introduce a trivial family-wise parameter condition for satisfying stochastic detailed balance. A reaction family *p* is said to satisfy condition (*D*_*p*_) if there exists a constant *ρ*_*p*_ > 0 such that

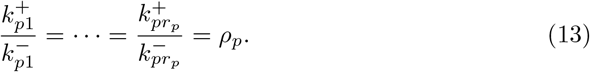

Under condition (*D*_*p*_), we have 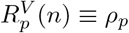. Consequently, if every reaction family *p* satisfies (*D*_*p*_), all parallelogram cycles are automatically balanced. Clearly, when there are no equivalent reactions, so that *r*_*p*_ = 1 for every reaction family *p*, condition (*D*_*p*_) holds automatically. This shows that the presence of equivalent reactions makes the problem more complicated.

We then discuss trivial parameter condition for linear-relation cycle. A linear-relation cycle associated with a nonzero integer vector *ξ* = (*ξ*_1_, …, *ξ*_*r*_) ∈ ℤ ^*r*^ satisfying Eq. (12) is balanced if and only if

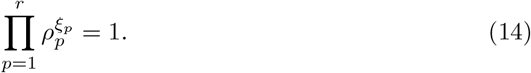

Therefore, conditions (*D*_*p*_) satisfied by all reaction families, together with condition (14) satisfied for every integer relation among the reaction vectors, give precisely the coefficient conditions for deterministic detailed balance [12, 15, 16].

Condition (*D*_*p*_) balances a reaction family through proportionality of its forward and backward rate constants. A different mechanism is provided by the following structural condition.

#### Definition 3 [16]

A reaction family *p* is said to satisfy the orthogonality condition (*O*_*p*_) if

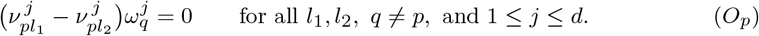

Condition (*O*_*p*_) means that every coordinate in which the source complexes of family *p* differ is left unchanged by all other reaction vectors. Its main consequence is the following Lemma, established in the proof of Theorem 3 in [16].

#### Lemma 1 [16]

*If reaction family p satisfies* (*O*_*p*_), *then*

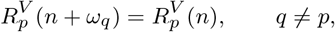

*whenever the two corresponding p-transitions are feasible*.

Combining Lemma 1 with condition (*D*_*p*_) gives the following sufficient condition for stochastic detailed balance.

#### Theorem 2

*[16] Suppose that the reaction vectors ω*_1_, …, *ω*_*r*_ *are linearly independent and that every reaction family p satisfies either* (*D*_*p*_) *or* (*O*_*p*_). *Then, for the given positive rate constants, the stochastic model satisfies stochastic detailed balance for every volume V* > 0.

Theorem 2 provides a mechanism for constructing systems that satisfy stochastic detailed balance without satisfying deterministic detailed balance. In particular, the orthogonality condition can replace the coefficient proportionality required by deterministic detailed balance. As a purely structural condition, it is independent of the rate constants and can generate systems whose global potentials exhibit a double-well structure [16]. The above discussion naturally leads to an interesting question: are there other structural or coefficient conditions that produce stochastic detailed balance? We will answer this question in the following sections.

## 3 Stochastic detailed balance can depend on volume

Before looking for additional mechanisms, one must decide whether stochastic detailed balance can be tested at a single volume. If it were independent of *V*, then the search would be considerably simpler. The next two examples show that stochastic detailed balance can indeed depend on the volume. The first example demonstrates this dependence through a linear-relation cycle, while the second demonstrates it through a parallelogram cycle.

### Example 1

Consider the following reaction system involving only one species *S*:

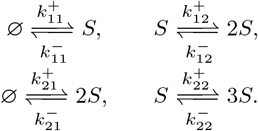

The distinct forward reaction vectors are

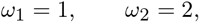

and hence 2*ω*_1_ − *ω*_2_ = 0. Clearly, we only need to consider this linearly dependent cycle. Choose arbitrary *a, b* > 0 and set

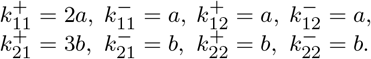

For the two reaction families, we have

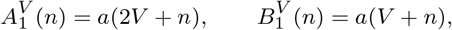

and

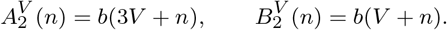

Therefore,

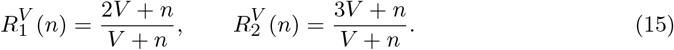

The relation 2*ω*_1_ − *ω*_2_ = 0 gives the feasible cycle

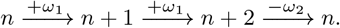

The left-hand side of the Kolmogorov cycle condition given in Eq. (9) along this cycle is

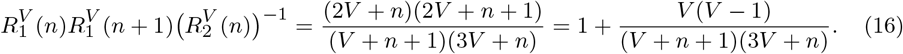

Clearly, this expression equals one if and only if *V* = 1. Thus the stochastic detailed balance is satisfied precisely when *V* = 1.

The first example shows that stochastic detailed balance can depend on the volume when the reaction vectors are linearly dependent. The next example shows that such volume dependence can also arise when the reaction vectors are linearly independent and hence no linear-relation cycles exist.

### Example 2

Consider a chemical system involving two species *S*_1_, *S*_2_ and two linearly independent reaction vectors

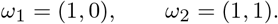

For family 1, take

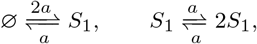

where *a* > 0. Direct calculation gives

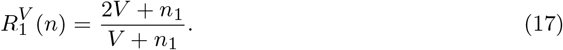

For family 2, take

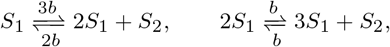

where *b* > 0. These two reactions both have reaction vector *ω*_2_ = (1, 1), and

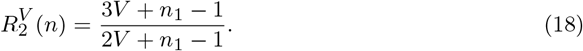

Let *n* = (*n*_1_, *n*_2_) be any state with *n*_1_ ≥ 2, so that every reaction channel is feasible along the parallelogram

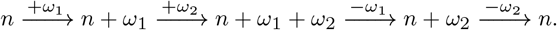

The left-hand side of Eq. (10) is given by

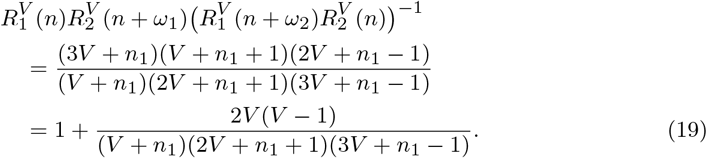

For *V* > 0 and *n*_1_ ≥ 2, the denominator is positive. Hence this cycle satisfies the Kolmogorov condition if and only if *V* = 1.

It remains to verify that every feasible cycle is balanced at *V* = 1. From Eqs. (17) and (18),

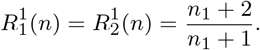

Both *ω*_1_ and *ω*_2_ increase *n*_1_ by one. Therefore, for every feasible parallelogram,

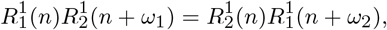

so all parallelogram cycles are balanced. Since *ω*_1_ and *ω*_2_ are linearly independent, there are no linear-relation cycles. Consequently, the chain satisfies stochastic detailed balanced exactly when *V* = 1.

These examples show that stochastic detailed balance requires precise relations between the rate constants and the volume. However, such tuning of parameters and volume has little practical relevance in real-world systems. This suggests that we need to distinguish whether satisfying stochastic detailed balance depends on the volume and parameters. Thus we give the following four types of stochastic detailed balance.

### Definition 4

Let *K* denote the set of all positive rate vectors. For a fixed volume *V*_0_ > 0 and a fixed rate vector *k*_0_ ∈ *K*, we introduce the following four types:

1. DB_11_(*V*_0_, *k*_0_) means that the stochastic model satisfies detailed balance at the fixed volume *V*_0_ and for the fixed rate vector *k*_0_.
2. DB_∞1_(*k*_0_) means that, for the fixed rate vector *k*_0_, the stochastic model satisfies detailed balance for every volume *V* > 0.
3. DB_1∞_(*V*_0_) means that, at the fixed volume *V*_0_, the stochastic model satisfies detailed balance for every positive rate vector *k* ∈ *K*.
4. DB_∞∞_ means that the stochastic model satisfies detailed balance for every volume *V* > 0 and every positive rate vector *k* ∈ *K*.

Clearly, in the notation of Definition 4, the first subscript of DB refers to the volume, while the second refers to the rate vector. Moreover, a subscript 1 ∞ indicates that the corresponding quantity is fixed, whereas a subscript ∞ indicates that detailed balance holds for every admissible choice of that quantity.

It is straightforward to see that the four types satisfy the following relation:

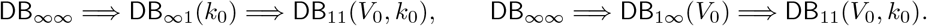

Moreover, Examples 1 and 2 show that

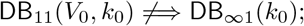

thus, for a fixed rate vector, stochastic detailed balance at one volume need not persist at other volumes. We will discuss the relationship among the four types in the next section.

## 4 Comparison between the four types of stochastic detailed balance

### 4.1 A simple condition for volume independence

We first introduce a trivial structural condition under which stochastic detailed balance at one volume automatically extends to every volume. This condition removes the volume dependence family by family and does not require the reaction vectors to be linearly independent.

#### Theorem 3

*Suppose that, for every reaction family p, all its source complexes have the same reaction order:*

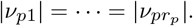

*Then* 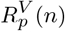 *is independent of V for every p. Consequently*,

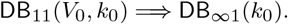

#### Proof

Fix a reaction family *p*, and denote the common reaction order of its source complexes by *m*_*p*_. From the definitions of 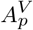 and 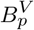,

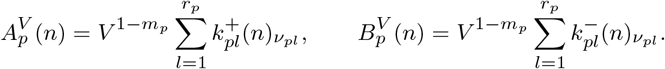

The common factor 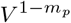 cancels, giving

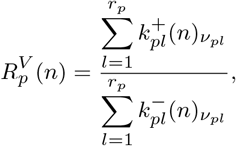

which is independent of *V*.

Now suppose that stochastic detailed balance holds at *V*_0_ for the fixed rate vector *k*_0_. For any feasible cycle, Eq. (9) gives

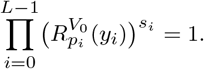

Finally, since 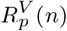 is independent of *V* for every reaction family *p*, detailed balance at *V*_0_ extends to every *V* > 0 immediately. Hence DB_∞1_(*k*_0_) holds. □

Although the condition in Theorem 3 is easy to verify, it is rather restrictive. For example, it is common and biologically important for an uncatalyzed reaction *S*_1_ ⇌ *S*_2_ to coexist with its enzyme-catalyzed counterpart *S*_1_ + *E* ⇌ *S*_2_ + *E*. These reactions have the same reaction vectors, but their source complexes have reaction orders one and two, respectively. To conclude, although Theorem 3 provides a convenient sufficient criterion for volume independence, it is too restrictive for a general comparison of the four types of stochastic detailed balance.

### 4.2 Detailed balance for arbitrary rate constants

We first discuss the case when the stochastic detailed balance is independent of the rate constants. The following theorem discusses the relationship between DB_1∞_(*V*_0_) and DB_∞∞_.

#### Theorem 4

*For every fixed V*_0_ > 0, *we have* DB_1∞_(*V*_0_) ⇐⇒ DB_∞∞_ ⇐⇒ [*ω*_1_, …, *ω*_*r*_ *are linearly independent and all* (*O*_*p*_) *hold*].

Clearly, Theorem 4 shows that DB_1∞_(*V*_0_) and DB_∞∞_ are actually equivalent, and it fully characterizes the network structures that satisfy DB_1∞_(*V*_0_). To prove Theorem 4, we need the following lemmas, which characterize the systems satisfying DB_1∞_(*V*_0_).

#### Lemma 2

*If the stochastic model satisfies stochastic detailed balance at a fixed volume V*_0_ *for every positive choice of rate constants, then ω*_1_, …, *ω*_*r*_ *are linearly independent*.

*Proof* Suppose, to the contrary, that the reaction vectors are linearly dependent. Since they have integer coordinates, there exists a nonzero vector *ξ* = (*ξ*_1_, …, *ξ*_*r*_) ∈ ℤ^*r*^ such that

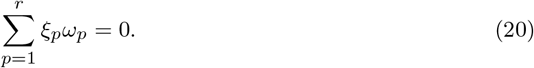

Starting from a state sufficiently far from the boundary, take *ξ*_*p*_ forward *p*-steps when *ξ*_*p*_ > 0 and −*ξ*_*p*_ backward *p*-steps when *ξ*_*p*_ *<* 0, in any fixed order. This gives a feasible cycle

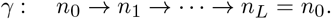

Using the notation introduced above, write

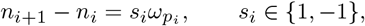

and let *y*_*i*_ be the lower endpoint of the *i*-th edge. By construction,

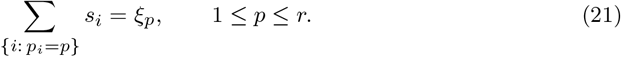

By Eq. (9), the Kolmogorov condition for *γ* is

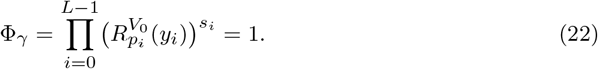

Choose *p*_0_ such that 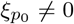, and replace

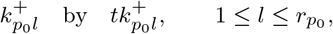

for some *t* > 0, leaving all other rate constants unchanged. Then 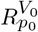 is multiplied by *t*, while every other 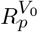 remains unchanged. Hence, by (21),

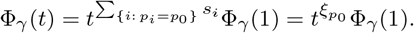

By assumption, stochastic detailed balance holds for every positive choice of rate constants.

Therefore,

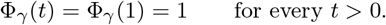

This gives

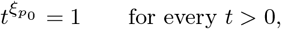

which is impossible because 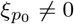. Thus *ω*_1_, …, *ω*_*r*_ are linearly independent. □

The above lemma shows that DB_1∞_(*V*_0_) implies the linear independence of the reaction vectors. The following lemma shows that DB_1∞_(*V*_0_) implies that every reaction family satisfies (*O*_*p*_).

#### Lemma 3

*If the stochastic model satisfies stochastic detailed balance at a fixed volume V*_0_ *for every positive choice of rate constants, then every reaction family satisfies* (*O*_*p*_).

#### Proof

Fix *p q* and choose *n* sufficiently large that the corresponding parallelogram is feasible. Choose the parameters of family *q* so that 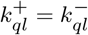 for every *l*. Then 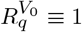, and Eq. (11)

Becomes

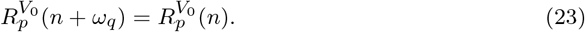

Note that the rate constants in family *p* remain arbitrary. Define

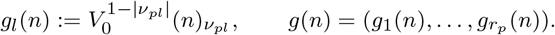

Writing 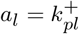 and 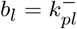, then Eq. (23) says

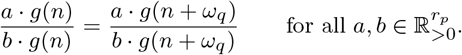

After cross multiplication, comparison of the coefficient of 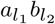 gives

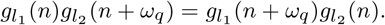

All quantities are positive for our feasible states, so division gives, for any *l*_1_, *l*_2_,

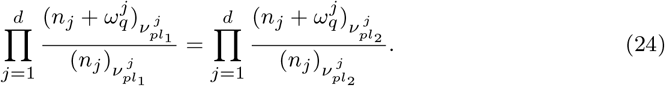

Now fix a coordinate *j* and fix all coordinates other than *n*_*j*_ at large values. Since Eq. (23) holds for every sufficiently large feasible state, Eq. (24) holds as an identity for all large integer values of *n*_*j*_. Move all factors independent of *n*_*j*_ to the right-hand side. We obtain

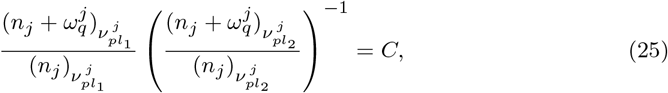

where *C* is independent of *n*_*j*_. Letting *n*_*j*_ → ∞ gives *C* = 1, because both falling-factorial quotients always tend to one for any parameters. Using

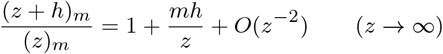

with *z* = *n*_*j*_,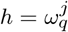, and 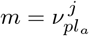 for *a* = 1, 2, the left-hand side of Eq. (25) is

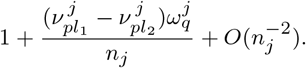

Since it is identically equal to 1 for all large *n*_*j*_, the coefficient of 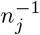 must vanish:

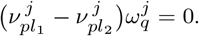

This holds for every *l*_1_, *l*_2_, every *q*≠= *p*, and every *j*; hence all (*O*_*p*_) hold. □

The preceding two lemmas establish the necessary structural conditions: if DB_1∞_(*V*_0_) holds, then the reaction vectors are linearly independent and every reaction family satisfies (*O*_*p*_). Since DB_∞∞_ implies DB_1∞_(*V*_0_), the same conditions are also necessary for DB_∞∞_. It remains only to prove that these structural conditions indeed imply both types.

#### Proof of Theorem 4

Suppose first that 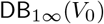 holds. It follows from Lemma 2 and Lemma 3 that in this case the reaction vectors *ω*_1_, …, *ω*_*r*_ are linearly independent and every reaction family satisfies (*O*_*p*_).

Conversely, suppose that *ω*_1_, …, *ω*_*r*_ are linearly independent and that (*O*_*p*_) holds for every *p*. It then follows from Theorem 2 that DB_∞∞_ holds. By definition, DB_∞∞_ implies DB_1∞_(*V*_0_) for every fixed *V*_0_ > 0. Therefore,

DB_1∞_(*V*_0_) ⇐⇒ DB_∞∞_ ⇐⇒ [*ω*_1_, …, *ω*_*r*_ are linearly independent and all (*O*_*p*_) hold].

We next provide an example to show that systems satisfying DB_∞∞_ indeed exist.

#### Example 3

Consider a system with four species *{S*_1_, *S*_2_, *E, F}* with the following reactions

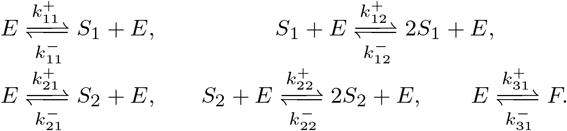

The reaction vectors are

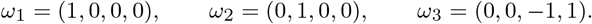

We now verify the orthogonality conditions. In family 1, the two source complexes *E* and *S*_1_ + *E* differ only in the first coordinate, which corresponds to species *S*_1_. Since both *ω*_2_ and *ω*_3_ have zero first coordinate, (*O*_1_) holds.

Similarly, in family 2, the source complexes *E* and *S*_2_ + *E* differ only in the second coordinate, which corresponds to species *S*_2_. Since both *ω*_1_ and *ω*_3_ have zero second coordinate, (*O*_2_) holds.

Family 3 has only one reversible pair, so (*O*_3_) is vacuous. The vectors *ω*_1_, *ω*_2_, *ω*_3_ are linearly independent. By Theorem 4, the model satisfies stochastic detailed balance for every volume *V* > 0 and every positive choice of the rate constants: DB_∞∞_.

This is nontrivial in the sense that deterministic detailed balance is not forced. Indeed, deterministic detailed balance would require

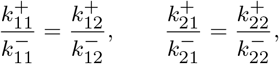

which need not hold for arbitrary parameters.

### 4.3 Stochastic detailed balance with fixed rate constants

We now consider stochastic detailed balance for a fixed set of rate constants. We show that, under suitable structural assumptions, stochastic detailed balance at a single volume extends to every volume, and that (*D*_*p*_) and (*O*_*p*_) are the only possible family-wise mechanisms underlying this property. Moreover, by violating such structural assumptions, we can construct more systems which satisfy stochastic detailed balance for any *V* with fixed rate constants. To formulate these assumptions, we first describe how one reaction family can affect another.

#### Definition 5

For two distinct reaction families *p* and *q*, we write

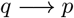

if there exist *l*_1_, *l*_2_ ∈ *{*1, …, *r*_*p*_*}* and *j* ∈ *{*1, …, *d}* such that

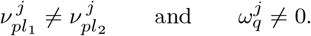

The relation *q* → *p* means that family *q* changes a coordinate in which the source complexes of family *p* differ. Therefore, a jump of family *q* may change the relative reaction rates within family *p*.

Accordingly, we write *q* ↛ *p* when *q* → *p* does not hold, and it can be characterized as

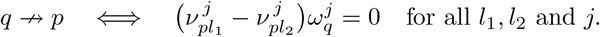

Moreover, it follows from the proof of Lemma 1 that we have

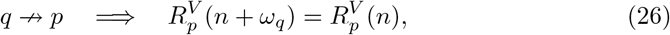

In particular, condition (*O*_*p*_) is equivalent to *q* ↛ *p* for every *q* ≠ *p*.

With the above definition and discussion, we now introduce two structural restrictions.

#### Definition 6 (Single-coordinate condition)

The network is said to satisfy condition (SC) if, for every reaction family *p*, there exists a coordinate *j*_*p*_ ∈ *{*1, …, *d}* such that

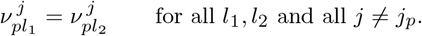

Thus the source complexes within each family may differ in at most one coordinate.

#### Definition 7 (No-bidirectional-influence condition)

The network is said to satisfy condition (NB) if influence between any two distinct reaction families cannot occur in both directions; that is,

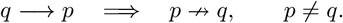

Condition (SC) restricts the differences among the source complexes of each family to a single coordinate, while condition (NB) prevents two families from affecting each other simultaneously. The following lemma shows their main use: when both conditions hold, stochastic detailed balance forces any family affected by another family to satisfy (*D*_*p*_).

#### Lemma 4

*Assume that conditions (SC) and (NB) hold and that* DB_11_(*V*_0_, *k*_0_) *holds. If q* → *p for some q*≠= *p, then the rate constants in family p satisfy* (*D*_*p*_).

To prove Lemma 4, we need the following lemma.

#### Lemma 5

*Let F be a rational function and let h*≠= 0. *If F* (*z* + *h*) = *F* (*z*) *for infinitely many distinct values of z at which both sides are defined, then F is constant*.

The proof of Lemma 5 is given in Appendix A. We now use it to prove Lemma 4.

#### Proof of Lemma 4

By condition (SC), there exists a coordinate *j*_*p*_ such that

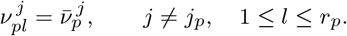

Thus the source complexes in family *p* may differ only in the *j*_*p*_-coordinate. Define

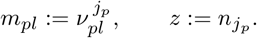

Here *m*_*pl*_ is the stoichiometric coefficient in the *j*_*p*_-coordinate of the source complex *v*_*pl*_.

Fix the remaining coordinates *n*_*j*_, *j*≠= *j*_*p*_, at sufficiently large values. We then have

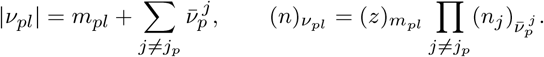

The factors common to all *l* cancel between 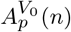 and 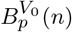. Therefore,

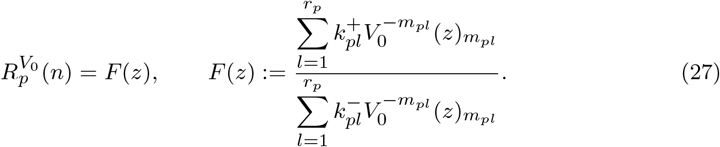

In particular, *F* is a rational function of the scalar variable *z*.

Since *q* → *p*, condition (NB) gives *p* ↛ *q*. Applying Eq. (26) with *p* and *q* interchanged, we obtain

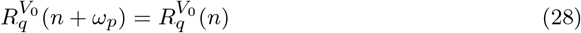

whenever the corresponding transitions are feasible. Moreover, DB_11_(*V*_0_, *k*_0_) implies the parallelogram condition

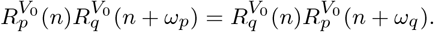

Using Eq. (28) and cancelling the positive factor 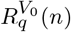, we obtain

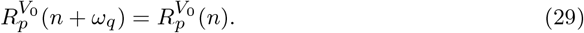

By the definition of *q* → *p*, family *q* changes a coordinate in which two source complexes of family *p* differ. Under condition (SC), this coordinate must be *j*_*p*_. Hence, for some *l*_1_, *l*_2_,

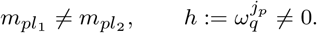

Since 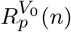 depends only on 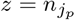, it follows from Eq. (27) that

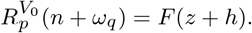

The fixed coordinates *n*_*j*_, *j* ≠ *j*_*p*_, can be chosen sufficiently large so that all required transitions are feasible for every sufficiently large integer *z*. By choosing infinitely many states sufficiently far from the boundary, Eq. (29) yields

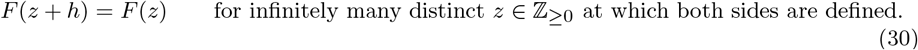

Lemma 5 therefore implies that *F* is constant. Since *h≠* 0, Lemma 5 implies that *F* is constant. Thus

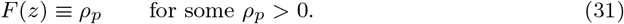

Substituting Eq. (31) into Eq. (27) and cross-multiplying yields

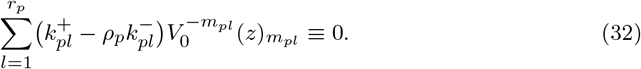

The integers 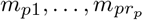 are distinct. Indeed, under condition (SC), two equal values of *m*_*pl*_ would give the same source complex. Since reactions in the same family have the same reaction vector, they would also have the same product complex and hence would be the same reaction.

The polynomials 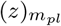 are monic and have distinct degrees, so they are linearly independent. It follows from Eq. (32) that

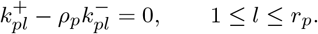

Consequently,

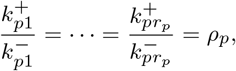

which is precisely condition (*D*_*p*_). □

We are now in a position to prove the following main result for stochastic detailed balance with fixed parameters.

#### Theorem 5

*Assume that the reaction vectors are linearly independent and that conditions* (SC) *and* (NB) *hold. For fixed positive rate constants k*_0_ *and any V*_0_ > 0, DB_11_(*V*_0_, *k*_0_) ⇐⇒ DB_∞1_(*k*_0_) ⇐⇒ [*every reaction family p satisfies either* (*D*_*p*_) *or* (*O*_*p*_)].

*Proof* Assume first that DB_11_(*V*_0_, *k*_0_) holds. Fix a reaction family *p*. If (*D*_*p*_) fails, Lemma 4 implies that

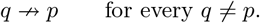

This is precisely condition (*O*_*p*_). Hence every reaction family *p* satisfies either (*D*_*p*_) or (*O*_*p*_).

Conversely, suppose that every reaction family satisfies either (*D*_*p*_) or (*O*_*p*_). Since the reaction vectors are linearly independent, Theorem 2 implies that the stochastic model satisfies detailed balance for every *V* > 0. Therefore, DB_∞1_(*k*_0_) holds. Finally,

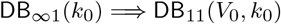

follows directly from the definitions. □

Theorem 5 shows that when the reaction vectors are linearly independent and conditions (SC) and (NB) both hold, stochastic detailed balance at a single volume implies stochastic detailed balance at every volume. Moreover, every reaction family *p* must satisfy either (*D*_*p*_) or (*O*_*p*_). Thus, (SC) and (NB) restrict stochastically detailed-balanced systems to these two structural types. We next demonstrate that violating either condition can give rise to new classes of systems satisfying stochastic detailed balance. Specifically, we construct two systems that satisfy stochastic detailed balance for every volume, although they contain reaction families satisfying neither (*D*_*p*_) nor (*O*_*p*_). The first example violates (SC) while satisfying (NB), whereas the second satisfies (SC) while violating (NB).

#### Example 4

Consider a chemical system involving *S*_1_ and *S*_2_ with two reaction vectors *ω*_1_ = (1, 0) and *ω*_2_ = (1, −1). Family 1 consists of

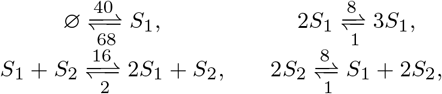

and family 2 is

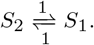

The sources 0, 2*S*_1_, *S*_1_ + *S*_2_, 2*S*_2_ of family 1 violate (SC), while (NB) holds because only family 2 influences family 1. Moreover,

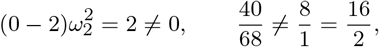

so (*O*_1_) and (*D*_1_) fail.

Then we have

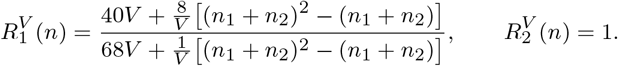

Since 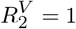, we have

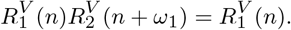

Moreover, *ω*_2_ = (1, −1) gives

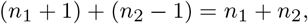

and 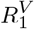 depends on *n* only through *n*_1_ + *n*_2_. Therefore

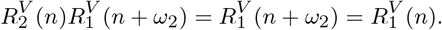

Then we have

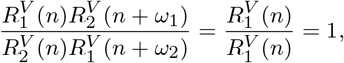

which satisfies the Kolmogorov cycle condition. Linear independence excludes linear-relation cycles; hence stochastic detailed balance holds for every *V* > 0.

For the deterministic system, we have

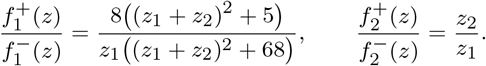

It follows from Theorem 1 that the equilibria satisfy

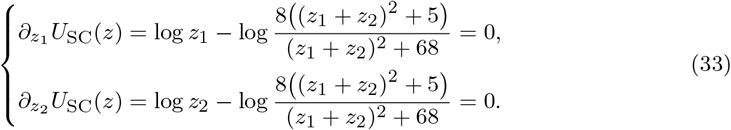

Solving Eq. (33) gives the three equilibria *{*(1, 1), (2, 2), (5, 5)*}*.

At (1, 1), the eigenvalues of the Hessian matrix are 1 and 2*/*9. Both are positive, so the Hessian matrix is positive definite and (1, 1) is a strict local minimum. At (2, 2), the eigenvalues of the Hessian matrix are 1*/*2 and −1*/*14. Since they have opposite signs, the Hessian matrix is indefinite and (2, 2) is a saddle point. At (5, 5), the eigenvalues of the Hessian matrix are 1*/*5 and 2*/*35. Both are positive, so the Hessian matrix is positive definite and (5, 5) is a strict local minimum. Therefore (1, 1) and (5, 5) are strict local minima, while (2, 2) is a saddle point. The resulting global potential is shown in the left panel of Fig. 1.

**Fig. 1.**
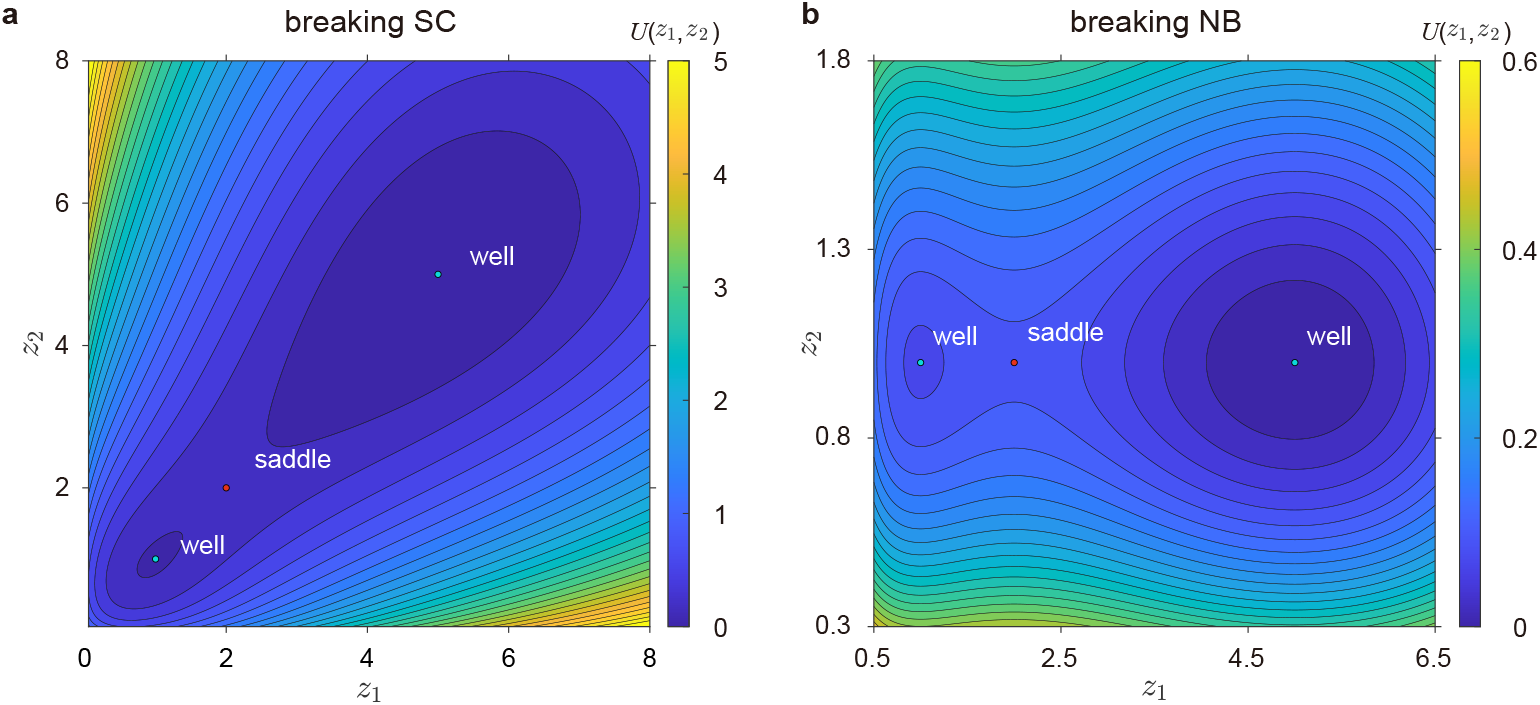
Contour plots of the global potentials generated by two different mechanisms. Panel (a) corresponds to Example 4, and panel (b) corresponds to Example 5. Each potential has two local minima separated by a saddle point.

#### Example 5

Let *ω*_1_ = (1, 0) and *ω*_2_ = (1, 1). Take

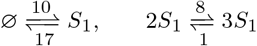

in family 1, and

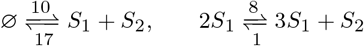

in family 2. Both families have sources 0, 2*S*_1_, so (SC) holds. Since both reaction vectors change the varying *S*_1_-coordinate, 1 → 2 and 2 → 1; hence (NB) fails. In each family,

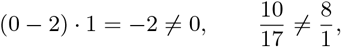

so both (*O*_*p*_) and both (*D*_*p*_) fail.

The stochastic rate ratios satisfy

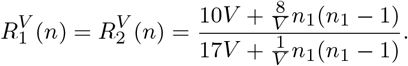

Here 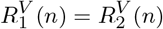. In addition, both *n* + *ω*_1_ and *n* + *ω*_2_ have first coordinate *n*_1_ + 1, and the displayed rate ratio depends only on the first coordinate. Hence

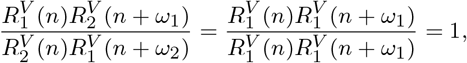

which satisfies the Kolmogorov cycle condition. Linear independence excludes linear-relation cycles, so stochastic detailed balance holds for every *V* > 0.

For the deterministic system, we have

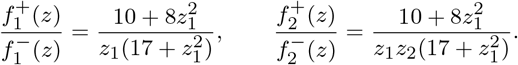

Similarly, using Theorem 1 we obtain that the equilibria satisfy

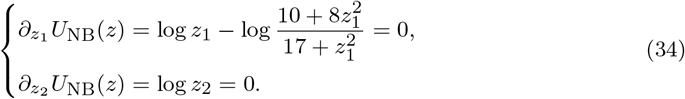

The second equation gives *z*_2_ = 1. Substituting this into the first equation gives the cubic equation

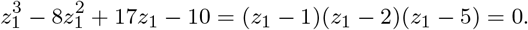

The stationary points of *U*_NB_ are (1, 1), (2, 1), and (5, 1).

At (1, 1), the eigenvalues of the Hessian matrix are 2*/*9 and 1. Both are positive, so the Hessian matrix is positive definite and (1, 1) is a strict local minimum. At (2, 1), the eigenvalues of the Hessian matrix are −1*/*14 and 1. Since they have opposite signs, the Hessian matrix is indefinite and (2, 1) is a saddle point. At (5, 1), the eigenvalues of the Hessian matrix are 2*/*35 and 1. Both are positive, so the Hessian matrix is positive definite and (5, 1) is a strict local minimum. The overall global potential is shown in the right panel of Fig. 1.

Thus far, the reaction vectors in all the examples presented above are linearly independent. We finally discuss the case where reaction vectors are linearly dependent. In this case, stochastic detailed balance requires both parallelogram cycles and linear-relation cycles to be balanced. Clearly, satisfying all parallelogram cycle conditions does not guarantee that the linear-relation cycle conditions are also satisfied. The presence of linear-relation cycles makes the characterization of stochastic detailed balance considerably more complicated. For systems with linear-relation cycles, whether stochastic detailed balance can arise without deterministic detailed balance is an interesting question for future research.

## 5 Conclusion

This paper systematically studies stochastic detailed balance in chemical reaction networks. Depending on whether stochastic detailed balance depends on the system volume and reaction rate parameters, we define four types of stochastic detailed balance and investigate their relationships. Our main results can be summarized as follows:

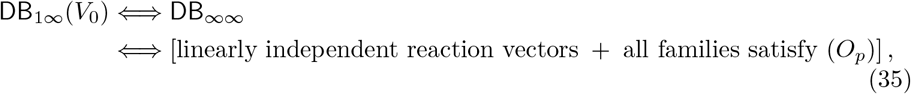

whereas, for one fixed parameter set,

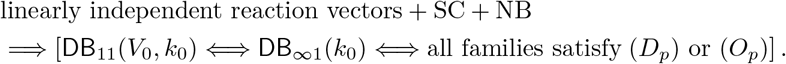

Moreover, using our results, we characterize systems that satisfy stochastic detailed balance for all volumes and parameter values, and construct new systems satisfying stochastic detailed balance by violating either (SC) or (NB).

We hope that the clarification of stochastic detailed-balance conditions and the construction of double-well systems in this work will help further illuminate the relationship between continuous deterministic models and discrete stochastic models, especially at the level of statistical physics.

## Acknowledgements

Youming Li acknowledges support from National Natural Science Foundation of China (No. 12401629), the Fundamental Research Funds for the Central Universities (No. ZYGX2024XJ043), and the Natural Science Foundation of Sichuan Province (No. 2026NSFSC0780).

## Author contributions

Shangbin Ma: Construction of counterexamples, Mathematical proofs, Writing–original draft, Programming; Youming Li: Conceptualization, Construction of counterexamples, Mathematical proofs, Supervision, Writing–review & editing.

## Data availability

No datasets were generated or analysed during the current study.

## Appendix A

### Proof of Lemma 5

Note that Lemma 5 states: Let *F* be a rational function and let *h /*= 0. If *F* (*z* + *h*) = *F* (*z*) for infinitely many values of *z* at which both sides are defined, then *F* is constant.

### Proof

Note that the difference

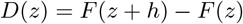

is a rational function, and we write it as

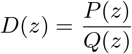

with *P, Q* polynomials and *Q* ≢ 0. At all points where the given equality holds and *D* is defined, we have *D*(*z*) = 0. Since there are infinitely many such points, the polynomial *P* has infinitely many zeros. Hence *P* ≡ 0, so *D* ≡ 0. Therefore,

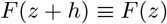

for every *z* at which both sides are defined.

Now write *F* = *P*_0_*/Q*_0_ in lowest terms. We claim that *F* has no finite poles. If *z*_0_ were a finite pole, then the identity *F* (*z* + *h*) ≡ *F* (*z*) would imply that *z*_0_ + *h* is also a pole; iterating, *z*_0_ + *kh* would be a pole for every integer *k*. These points are all distinct because *h*≠ 0. This gives infinitely many finite poles, which is impossible for a rational function. Thus *F* has no finite poles.

Since *F* = *P*_0_*/Q*_0_ is in lowest terms and has no finite poles, *Q*_0_ has no zeros. Over the complex numbers a nonconstant polynomial at least has a zero, so *Q*_0_ must be constant. Hence *F* is a polynomial.

Finally, a polynomial invariant under a nonzero shift must be constant. Indeed, suppose that *F* has degree *d* ≥ 1 with leading term *cz*^*d*^, where *c*≠ 0. Then *F* (*z* + *h*) − *F* (*z*) has leading term

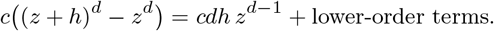

This is nonzero because *c*≠ 0, *d* ≥ 1, and *h*≠ 0, contradicting *F* (*z* + *h*) ≡ *F* (*z*). Therefore *F* has degree 0, so *F* is constant.

